# Septate junction proteins are required for egg elongation and border cell migration during oogenesis in Drosophila

**DOI:** 10.1101/2020.10.19.346064

**Authors:** Haifa Alhadyian, Dania Shoaib, Robert E. Ward

## Abstract

Protein components of the invertebrate occluding junction - known as the septate junction (SJ) - are required for morphogenetic developmental events during embryogenesis in *Drosophila melanogaster*. In order to determine whether SJ proteins are similarly required for morphogenesis during other developmental stages, we investigated the localization and requirement of four representative SJ proteins during oogenesis: Contactin, Macroglobulin complement-related, Neurexin IV, and Coracle. A number of morphogenetic processes occur during oogenesis, including egg elongation, formation of dorsal appendages, and border cell migration. We found that all four SJ proteins are expressed in egg chambers throughout oogenesis, with the highest and most sustained levels in the follicular epithelium (FE). In the FE, SJ proteins localize along the lateral membrane during early and mid-oogenesis, but become enriched in an apical-lateral domain (the presumptive SJ) by stage 10b. SJ protein relocalization requires the expression of other SJ proteins, as well as rab5 and rab11 in a manner similar to SJ biogenesis in the embryo. Knocking down the expression of these SJ proteins in follicle cells throughout oogenesis results in egg elongation defects and abnormal dorsal appendages. Similarly, reducing the expression of SJ genes in the border cell cluster results in border cell migration defects. Together, these results demonstrate an essential requirement for SJ genes in morphogenesis during oogenesis, and suggests that SJ proteins may have conserved functions in epithelial morphogenesis across developmental stages.

**Article Summary:** Septate junction (SJ) proteins are essential for forming an occluding junction in epithelial tissues of *Drosophila melanogaster*, and also for morphogenetic events that occur prior to the formation of the junction during embryogenesis. Here we show that SJ proteins are expressed in the follicular epithelium of egg chambers during oogenesis and are required for morphogenetic events including egg elongation, dorsal appendages formation, and border cell migration. Additionally, the formation of SJs during oogenesis is similar to that in embryonic epithelia.

## Introduction

The septate junction (hereafter referred to as SJ) provides an essential paracellular barrier to epithelial tissues in invertebrate animals (Noirot-timothée *et al.* 1978). As such, the SJ is functionally equivalent to the tight junction in vertebrate tissues, although the molecular components and ultrastructure of these junctions differ (reviewed in Izumi and Furuse 2014). Studies in *Drosophila* have identified more than 20 proteins that are required for the organization or maintenance of the SJ (Fehon *et al.* 1994; Baumgartner *et al.* 1996; Behr *et al.* 2003; Paul *et al.* 2003; Genova and Fehon 2003; Faivre-Sarrailh *et al.* 2004; Wu *et al.* 2004; Wu *et al.* 2007; Tiklová *et al.* 2010; Nelson *et al.* 2010; Ile *et al.* 2012; Bätz *et al.* 2014; Hall *et al.* 2014). Given that some of these genes have clear developmental functions (e.g. *coracle’s* name derives from its dorsal open embryonic phenotype; (Fehon *et al.* 1994), we previously undertook an examination of the developmental requirements for a set of core SJ genes (Hall and Ward 2016). We found that all of the genes we analyzed (9 in all) are required for morphogenetic developmental events during embryogenesis including head involution, dorsal closure and salivary gland organogenesis. Interestingly, these embryonic developmental events occur prior to the formation of an intact SJ, suggesting that these proteins have a function independent of their role in creating the occluding junction (Hall and Ward 2016). Since strong loss of function mutations in every SJ gene are embryonic lethal (due to these morphogenetic defects and/or a failure in establishing a blood-brain barrier in glial cells; Baumgartner *et al.* 1996), only a few studies have examined the role of SJ proteins in morphogenesis at a later stages of development. These studies have revealed roles for SJ proteins in planar polarization of the wing imaginal disc, for epithelial rotations in the eye and genital imaginal discs, and ommatidia integrity (Lamb *et al.* 1998; Venema *et al.* 2004; Moyer and Jacobs 2008; Banerjee *et al.* 2008).

To further explore the role of SJ proteins in morphogenesis beyond the embryonic stage, we set out to examine the expression and function of a subset of SJ genes in the *Drosophila* egg chamber during oogenesis. Each of the two *Drosophila* ovaries is comprised of approximately 16-20 ovarioles, which are organized into strings of progressively developing egg chambers (Figure 1A). Each egg chamber forms in a structure called the germarium, where the germline and somatic stem cells reside. Once the egg chamber is formed, it leaves the germarium as 16-cell germline cyst consisting of 15 nurse cells and an oocyte surrounded by a layer of somatic follicle cells (FCs) (Figure 1B). An egg chamber undergoes 14 developmental stages ending in a mature egg that is ready for fertilization (reviewed in Horne-Badovinac and Bilder 2005). Interfollicular cells called stalk cells connect egg chambers to each other. During oogenesis, the follicular epithelium (FE) undergoes several morphogenetic events including border cell migration, dorsal appendage formation and egg elongation (reviewed in Horne-Badovinac and Bilder 2005; reviewed in Duhart *et al.* 2017).

**Figure 1.**
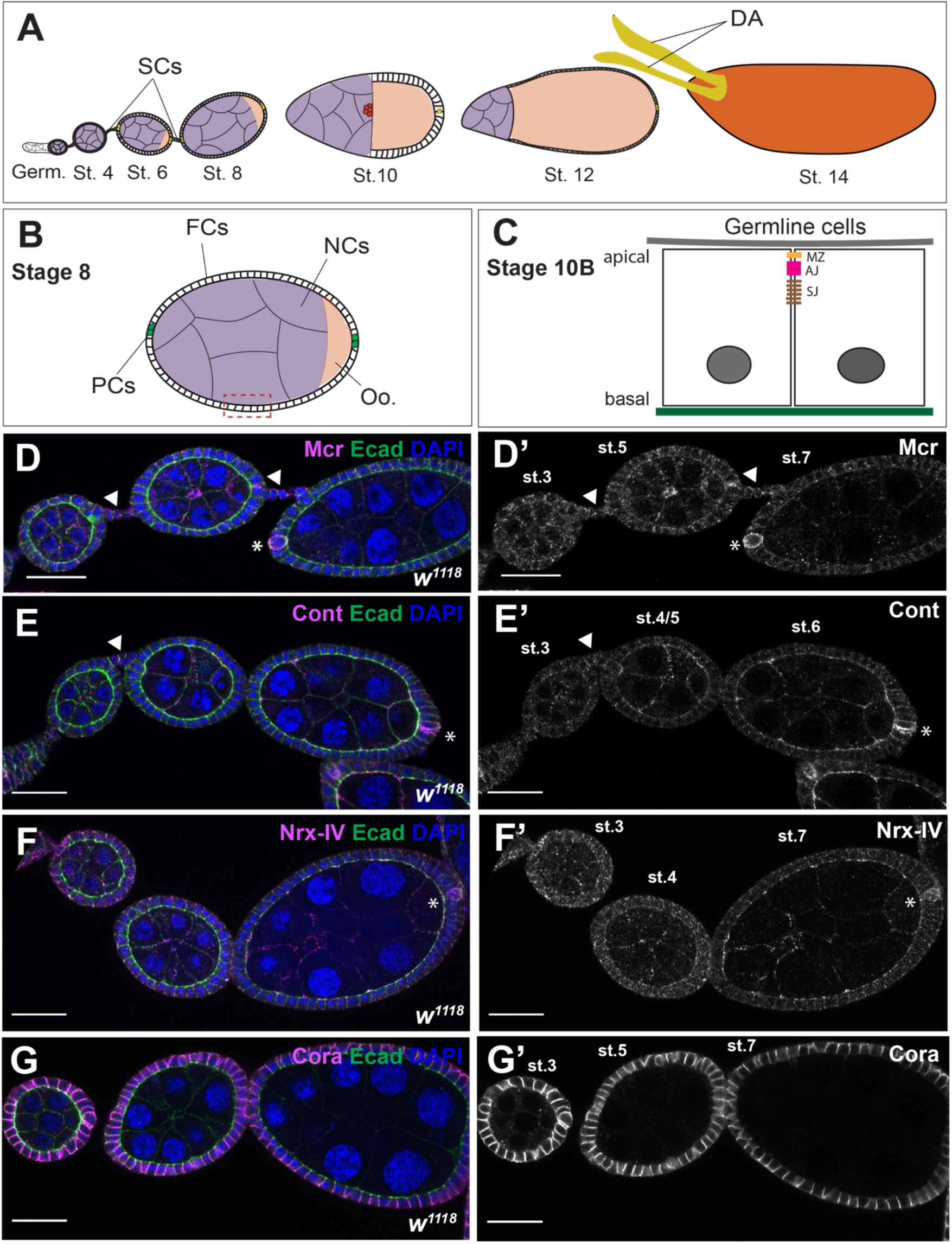
Mcr, Cont, Nrx-IV, and Cora expression during early stages of oogenesis. (A) Diagram of a female *Drosophila* ovariole. Egg chambers are formed in the most anterior region of the ovariole called the germarium (Germ). Each egg chamber undergoes 14 developmental stages while connected to each other through stalk cells (SCs) to form a mature stage 14 egg. (B) Diagram of a stage 8 egg chamber. The egg chamber consists of 15 nurse cells (NCs) and one oocyte (Oo.), which are surrounded by a monolayer of follicle cells (FCs). At the anterior and posterior ends of an egg chamber resides a pair of differentiated FCs called polar cells (PCs). (C) Diagram of a lateral view of a portion of a stage 10B egg chamber. FCs face the germline and have defined apical-basal polarity with the apical surface facing the germline and a lateral junctional complex consisting of a marginal zone (MZ), an adherens junction (AJ), and a septate junction (SJ). (D-G) Confocal optical sections of wild type early stages egg chambers stained with antibodies against Mcr (D), Cont (E), Nrx-IV (F), and Cora (G), and co-stained with antibodies against Ecad (green) and labeled with DAPI. All four SJ proteins are expressed throughout the egg chamber along FC membranes, including SCs (arrowheads in B and D, and data not shown for Nrx-IV and Cora) and in the NCs. Mcr, Cont, and Nrx-IV (D-F) are found along the membrane and in puncta, whereas Cora is found predominantly at the membrane (G). In addition, Mcr, Cont, and Nrx-IV are highly expressed in the PCs (asterisks in D-F), whereas Cora is expressed in these cells with same level of expression relative to the FCs (data not shown). Note that the focal plane of these images shows strong staining in PCs in only one side of the egg chamber, but Mcr, Cont and Nrx-IV are equally expressed in both anterior and posterior PCs. Anterior is to the left in each ovariole. Scale bar = 25μm.

Previous studies have revealed that a few core components of the SJ are expressed in the ovary, including Macroglobulin complement-related (Mcr), Neurexin IV (Nrx-IV), Contactin (Cont), Neuroglian (Nrg), and Coracle (Cora) (Wei *et al.* 2004; Schneider *et al.* 2006; Maimon *et al.* 2014; Hall *et al.* 2014; Ben-Zvi and Volk 2019), although the developmental expression pattern and subcellular localizations of these proteins have not been thoroughly investigated. Furthermore, ultrastructural analysis has revealed the presence of mature SJs in the FE by stage 10/10B of oogenesis (Figure 1C), while incipient SJ structures have been observed in egg chambers as early as stage 6 (Mahowald 1972; Müller 2000). The biogenesis of SJs in embryonic epithelia is a multistep process in which SJ proteins are initially localized along the lateral membrane, but become restricted to an apical-lateral region (the SJ) in a process that required endocytosis and recycling of SJ proteins (Tiklová *et al.* 2010). How SJ maturation occurs in the FE is unknown.

Here, we analyzed the expression and subcellular localization of the core SJ proteins Mcr, Cont, Nrx-IV, and Cora throughout oogenesis. We find that all of these SJ proteins are expressed in the FE throughout oogenesis. Interestingly, Mcr, Cont, Nrx-IV, and Cora become enriched at the most apical-lateral region of the membrane in stage 10b/11 egg chambers, coincident with the formation of the SJ revealed by electron microscopy (Mahowald 1972; Müller 2000). Similar to the biogenesis of SJs in the embryo, this enrichment of SJ proteins to the presumptive SJ requires the function of other SJ genes, as well as *Rab5* and *Rab11.* Functional studies using RNA interference (RNAi) of SJ genes in FCs results in defects in egg elongation, dorsal appendage morphogenesis and border cell migration. Together, these results reveal a strong similarity in the biogenesis of SJ between embryonic and follicular epithelia, demonstrate that at least some components of the SJs are required for morphogenesis in the ovary, and suggest that these roles may be independent of their role in forming an occluding junction.

## Material and methods

### Fly stocks

All *Drosophila* stocks were maintained on media consisting of corn meal, sugar, yeast, and agar on shelves at room temperature or in incubators maintained at a constant temperature of 25°C. GAL4 lines used in this study are as follows: *GR1-GAL4* (Bloomington Drosophila Stock Center (BDSC) #36287), *Slbo-GAL4, UAS-mCD8-GFP* (BDSC#76363), and *C306-GAL4; GAL80^ts^/Cyo* (a gift from Jocelyn McDonald, Kansas State University, Manhattan, Kansas). RNAi stocks used for these studies are as follows: *UAS-Mcr-RNAi* (BDSC#65896 and Vienna Drosophila Resources Center (VDRC)#100197), *UAS-Cora-RNAi* (BDSC#28933 and VDRC#9787), *UAS-Nrx-IV-RNAi* (BDSC#32424 and VDRC#9039), *UAS-Cont-RNAi* (BDSC#28923), *UAS-mCherry-RNAi* (BDSC#35787), *UAS-Lac-RNAi* (BDSC#28940), and *UAS-Sinu-RNAi* (VDRC#44929). *UAS-Rab5^DN^* (BDSC#9771) was used to inhibit normal Rab5 function and *UAS-Rab11-RNAi* (BDSC#27730) was used to knock down Rab11 in the follicle cells. *UAS-GAL80^ts^* (BDSC#7108) was used to conditionally inhibit *GR1-GAL4* activity in the *UAS-Rab11-RNAi* experiment. *UAS-GFP* (BDSC#1521) was crossed to *GR1-GAL4* as a control for the egg shape experiments. *Slbo-GAL4, UAS-mCD8-GFP* was crossed to *UAS-mCherry-RNAi* as a control for one set of border cell migration studies, whereas *C306-GAL4; GAL80^ts^/Cyo* was crossed to *UAS-Dcr* (BDSC#24646) as a control for the other set of border cell migration studies. *w^1118^* (BDSC# 5905) was used as the wild type stock for determining the expression of Mcr, Cont, Nrx-IV and Cora in the follicle cells.

### Fly staging

*w^1118^* 1-2-day-old females and males were collected and reared at 25°C on fresh food sprinkled with yeast for five to six days before the females were dissection for antibody staining. For egg elongation analyses, crosses were maintained at 25°C, and 1-2-day-old females (control and *UAS-RNAi*-expressing) were mated with sibling males and maintained at 29-30°C for 3 days before dissection. For border cell migration analyses, *Slbo-GAL4* crosses were kept at 25°C, whereas *C306-GAL4/UAS-Dcr; GAL80^ts^/SJ-RNAi* crosses were kept at 18°C to prevent GAL4 activation. 1-2-day-old flies with the appropriate genotype (*Slbo-GAL4, UAS-mCD8-GFP/UAS-RNAi* or *C306-GAL4/UAS-Dcr;UAS-RNAi;GAL80^ts^*) were shifted to 29-30°C for 48 hours before dissection. It should be noted that by crossing *UAS-GFP* to *C306-GAL4*, we observed the expression of GFP in polar cells in stage 10, but not stage 9 egg chambers (data not shown). For *Rab11-RNAi* experiment, crosses were maintained at 18°C and 2-3-day-old males and females with the appropriate genotype (*GR1-GAL4>UAS-mCherry-RNAi, UAS-GAL80ts or GR1-GAL4>UAS-Rab11-RNAi, UAS-GAL80ts*) were collected and reared at 29°C-30°C overnight before dissection. For the *Rab5^DN^* experiment, crosses were maintained at 25°C, and 1-2-day-old females were mated to sibling males and maintained at 29-30°C for 3 days before dissection.

### Egg aspect ratio measurements

Stage 14 egg chambers were selected for analysis based on the overall morphology of the egg and the absence of nurse cells nuclei by DAPI staining. Stage 14 egg chambers that have irregular edges or touch other egg chambers were excluded from the analysis to prevent inaccurate measurements. Egg length (anterior-posterior) and width (dorsal-ventral) were measured using the ImageJ/Fiji (http://fiji.sc) (Schindelin *et al.* 2012) straight-line tool, and aspect ratio was calculated as length divided by width using Excel Microsoft.

### Border cell migration quantification

Stage 10 egg chambers were identified based on the morphology of the egg (oocyte occupies half the egg chamber, whereas the other half is occupied by the nurse cells and centripetal cells). We used the GFP signal in Slbo-GAL4 crosses and DAPI and/or Fas3 staining in c306-GAL4 crosses to identify the location of the border cell cluster in stage 10 egg chambers. The location of the border cell cluster was quantified and grouped into four categories - complete, incomplete, failed migration, and disassociated cluster based on the location of the cluster relative to the oocyte in a stage 10 egg chamber (Figure 6). In some cases, border cell clusters display two phenotypes such as complete and dissociated. In this case, we quantified both phenotypes in one egg chamber.

### Immunostaining and image acquisition

Ovaries were dissected in 1X Phosphate-buffered saline (PBS), fixed in 4% Paraformaldehyde for 20 minutes, washed three times in 1X PBS, and then permeabilized in a block solution (1X PBS + 0.1% Triton + 1% Normal Donkey Serum) for 30 minutes before incubation with primary antibodies either overnight at 4°C or 2-4 hours at room temperature (~23-25°C). The following antibodies were used at the given dilutions: guinea pig (gp) anti-Cont 1:2000 (*Faivre-Sarrailh et al.* 2004) and rabbit (rab) anti-Nrx-IV 1:500 (Baumgartner *et al.* 1996) obtained from Manzoor Bhat, University of Texas Health Science Center, San Antonio, TX, gp anti-Mcr 1:1000 (Hall *et al.* 2014), mouse (m) anti-Cora (C566.9 and C615.16 mixed 1:1, obtained from the Developmental Studies Hybridoma Bank (DSHB) at the University of Iowa, Iowa City, IA; Fehon *et al.* 1994) 1:50, rat anti-*D*E-cad (DCAD2, DSHB) 1:27, and m anti-Fas3 (7G10, DSHB) 1:260. DAPI (1mg/ml) was used at a dilution of 1:1000. Secondary antibodies were obtained from Jackson ImmunoResearch Laboratories (West Grove, Pennsylvania, USA) and were used at 1:500.

Images were acquired using an Olympus FV1000 confocal microscope equipped with Fluoview software (version 4.0.3.4). Objectives used included an UPLSAPO 20X Oil (NA:0.85), a PLAPON 60X Oil (NA: 1.42), and an UPLSAPO 100X Oil (NA:1.40). Stage 14 egg chambers were imaged using Nikon Eclipse 80*i* compound microscope. Raw images were rotated and cropped in ImageJ/Fiji. Micrographs were adjusted for brightness using Adobe Photoshop 21.1.1 (San Jose, CA) or Image/Fiji. Adobe Illustrator 24.1 was used to compile the figures.

### Statistical Analysis

An unpaired *t*-test was used to calculate the P values in egg chamber aspect ratio between control and SJ mutant stage 14 egg chambers using GraphPad Prism 8 (https://www.graphpad.com) (version 8.4.2).

### Data Availability

Fly stocks are available upon request. Supplemental files are available at FigShare. Figure S1 shows the efficiency of RNAi knock-down in the FE of stage 12 egg chambers. Figure S2 shows the range of dorsal appendage phenotypes found in *GR1>SJ-RNAi* stage 14 egg chambers. Figure S3 shows the expression of Contactin during border cell migration. Figure S4 shows the expression of Nrx-IV during border cell migration. Figure S5 shows the expression of Coracle during border cell migration. The authors affirm that all the data necessary for confirming the conclusions of the article are present within the article and figures.

## Results

### Septate junction proteins are expressed in follicle cells throughout oogenesis

While a few SJ *proteins* have previously been reported to be expressed in the *Drosophila* ovary (Wei *et al.* 2004; Schneider *et al.* 2006; Hall *et al.* 2014; Maimon *et al.* 2014; Felix *et al.* 2015; Ben-Zvi and Volk 2019), a thorough analysis of their tissue distribution and subcellular localization throughout oogenesis is lacking. We therefore examined the spatial and temporal expression of four SJ proteins: Mcr, Cont, Nrx-IV and Cora (Fehon *et al.* 1994; Baumgartner *et al.* 1996; Faivre-Sarrailh *et al.* 2004; Bätz *et al.* 2014; Hall *et al.* 2014). These four proteins are core components of the junction for which well-characterized antibodies are available.

At early stages of oogenesis (stages 2-8), Mcr, Cont, and Nrx-IV all localize in puncta at the lateral membrane of FCs and nurse cells, and show a punctate distribution in these cells (Figure 1D-F). Mcr, Cont, and Nrx-IV are also more strongly expressed in polar cells (PCs) than the surrounding FCs (asterisks in Figure 1D-F). Cora is more uniformly localized along the lateral membrane of the FCs, including the PCs (Figure 1G and data not shown). These SJ proteins are additionally expressed in stalk cells (arrowheads in Figure 1 D and E and data not shown). Beginning at stage 10B, Mcr, Nrx-IV, Cont and Cora are gradually enriched at the apical-lateral membrane of the FCs just basal to the AJ. This localization is complete by stage 11 and persists to the end of oogenesis (arrows in Figure 2B, D, F, and H). The timing of this apical-lateral enrichment of Mcr, Cont, Nrx-IV and Cora coincides with the maturation of the SJ in the FCs based upon ultrastructural analysis (Mahowald 1972; Müller 2000), and so we will refer to this region as the presumptive SJ. Finally, all of these SJ proteins continue to be expressed in the FCs until stage 14 of oogenesis (Figure 2C, E, G and I).

**Figure 2.**
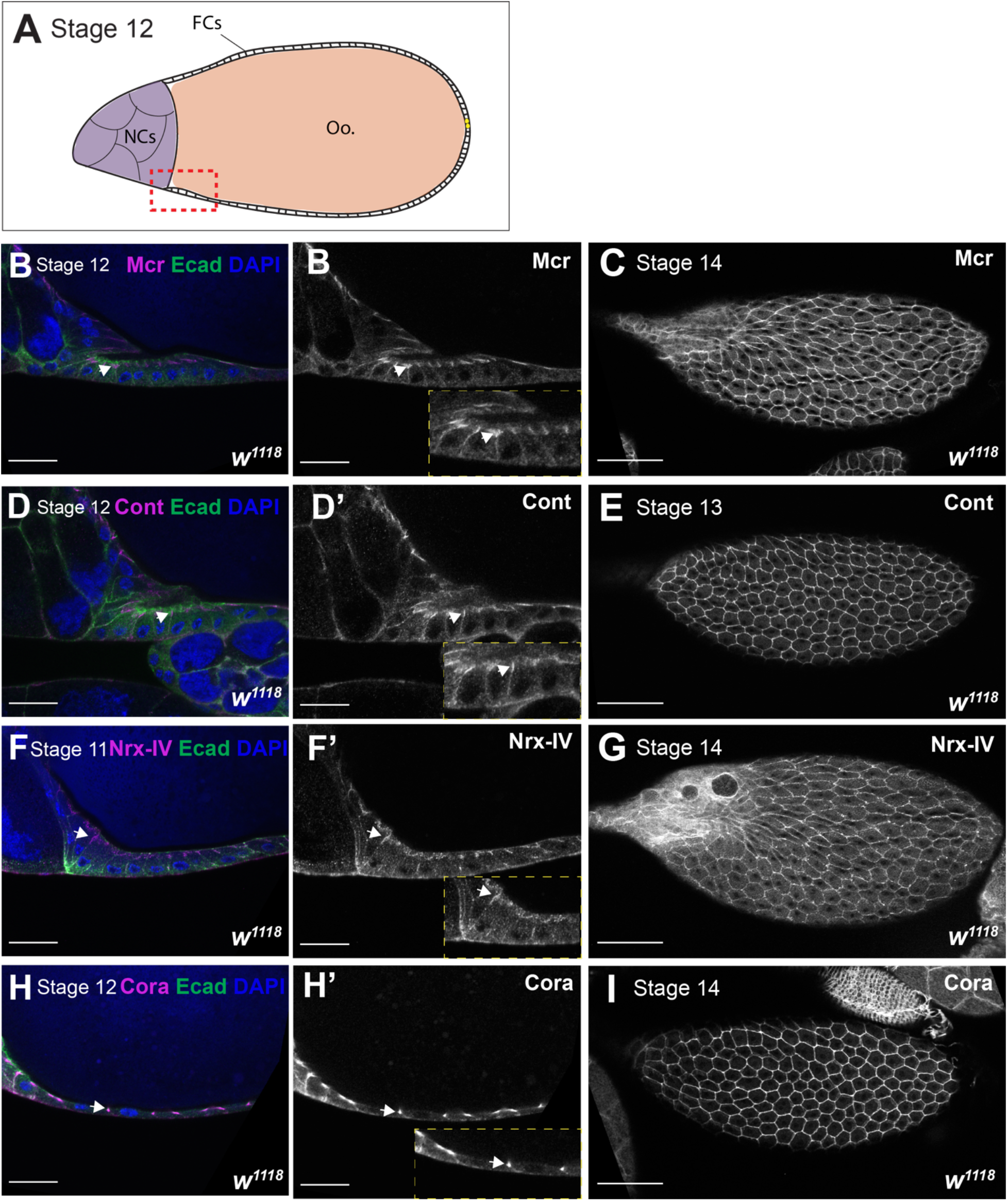
Mcr, Cont, Nrx-IV, and Cora localization at later stages of oogenesis. Confocal optical sections of wild type stage 11 – 12 egg chambers (B, D, F, and H) or stage 14 (C, E, G, and I) stained with antibodies against Mcr (B and C), Cont (D and E), Nrx-IV (F and G), or Cora (H and I) and co-stained with antibodies against Ecad (green) and labeled with DAPI (blue). Individual channels for SJ protein staining are shown in (B’-H’). The location of these sections is shown on the stage 12 egg chamber in (A), and is overlaying the boundary between the oocyte (Oo.) and nurse cells (NCs). Note that Mcr, Cont, Nrx-IV, and Cora become enriched at the apical-lateral region of FCs membrane (arrow in B, D, F, and H). The expression of all of these proteins persists in stage 14 egg chambers (C, E, G, and I). Anterior is to the left. Scale bar = 25μm in B, D, F, and H and 100μm in C, E, G, and I.

### SJ proteins are required for egg elongation and dorsal appendage morphogenesis

Given our findings that these four SJ proteins are expressed in the FE throughout oogenesis, and our previous studies indicating a role for SJ proteins in morphogenesis, we wondered whether SJ proteins might be required for morphogenetic processes in the FE. The FE plays critical roles in shaping the egg chamber and producing the dorsal appendage, while a subset of FE cells participates in border cell migration to form the micropyle (Montell 2003; Horne-Badovinac 2020). Because SJ mutant animals die during embryogenesis, we used the *GAL4 UAS-RNAi* system to knock-down the expression of SJ proteins in the FCs (Brand and Perrimon 1993). To knock down expression of SJ proteins throughout the majority of oogenesis, we used *GR1-GAL4*, which is expressed in the FCs from stage 4 to 14 of oogenesis (Gupta and Schüpbach 2003; Wittes and Schüpbach 2018). In all, we tested Bloomington Transgenic RNAi Project (TRiP) lines made against six different SJ genes (*Cont, cora, Mcr, lac, Nrx-IV*, and *sinu).* To examine overall egg chamber shape, we dissected stage 14 egg chambers from females expressing *SJ-RNAi* under the control of *GR1-GAL4*, imaged them on a compound microscope, and determined the aspect ratio of the egg chambers using measurements of egg chamber length and width using ImageJ/Fiji. Control stage 14 egg chambers (*GR1-GAL4>UAS-GFP*) had a mean aspect ratio of 2.3 (Figure 3A and J). In contrast, the aspect ratio of stage 14 egg chambers from all *GR1-GAL4>SJ-RNAi* is statistically significant than control egg chambers (aspect ratios from 1.7 to 2.1; Figure 3B-G and J). All *SJ-RNAi* egg chambers are significantly shorter than control (Figure 3H), and all but *Mcr-RNAi* and *Cont-RNAi* are also wider than control egg chamber (Figure 3I). To address the specificity of these results we also tested VDRC RNAi lines directed against Mcr, Cora, and Nrx-IV, and found similar effects on egg shape (Figure 3). We also examined SJ protein expression in late-stage egg chambers for Mcr, Cora, and Nrx-IV-RNAi to demonstrate that the RNAi was efficiently knocking down protein expression in this tissue (Figure S1).

**Figure 3.**
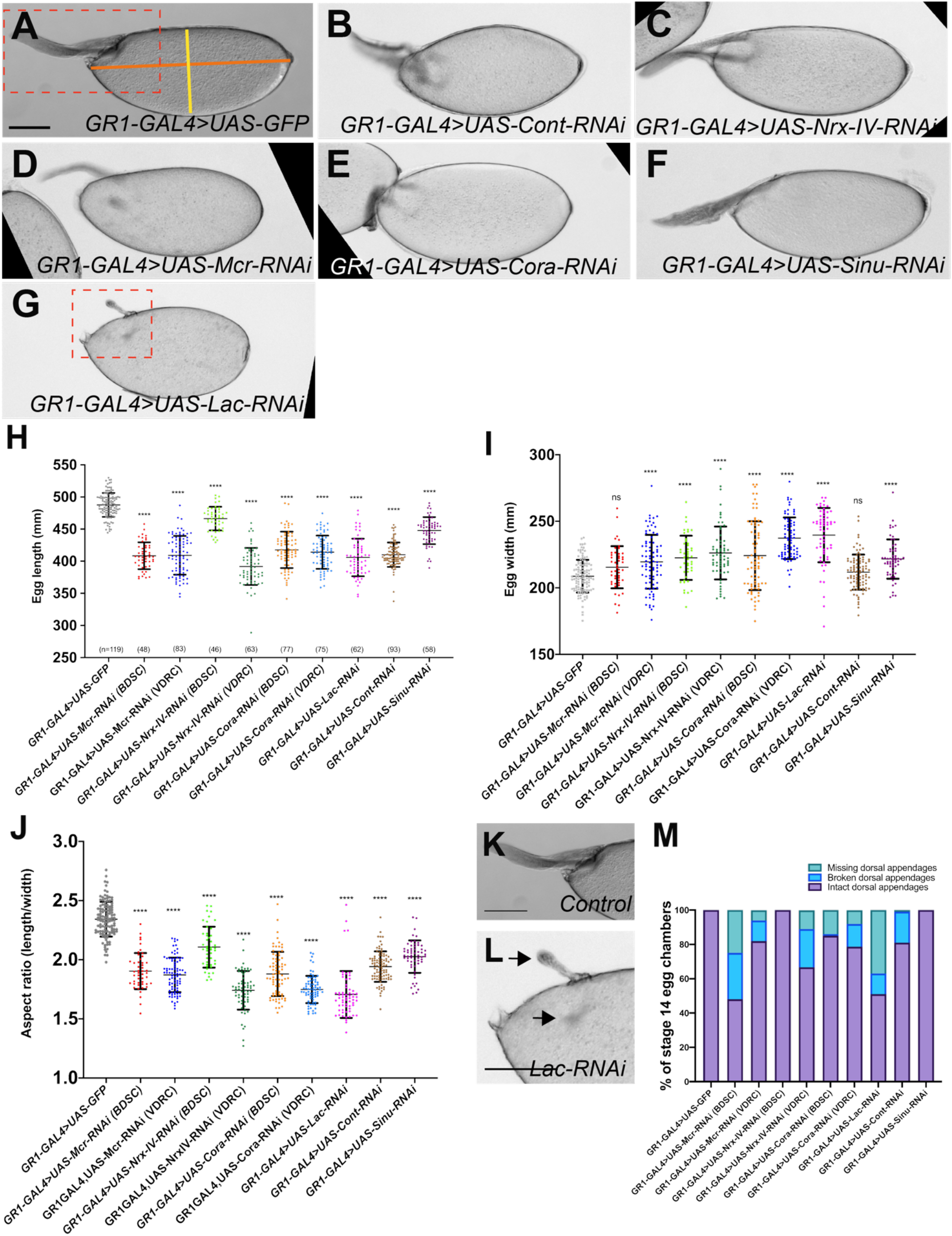
*SJ* genes are required for egg elongation. (A-G) Brightfield photomicrographs of stage 14 egg chambers. (A) *GR1-GAL4>UAS-GFP*, (B) *GR1-GAL4>UAS-Cont-RNAi*, (C) *GR1-GAL4>UAS-Nrx-IV-RNAi*, (D) *GR1-GAL4>UAS-Mcr-RNAi*, (E) *GR1-GAL4>UAS-Cora-RNAi*, (F) *GR1-GAL4>UAS-Sinu-RNAi*, and (G) *GR1-GAL4>UAS-Lac-RNAi.* Images in this figure represent average phenotypes observed in each genotype. (H and I) Quantification of length and width of stage 14 egg chambers from control and *SJ-RNAi* egg chambers. (J) Quantification of the aspect ratio of length (orange line in A) to width (yellow line in A) from control and *SJ-RNAi* stage 14 egg chambers. Note that the aspect ratio of all *SJ-RNAi* expressing egg chambers are statistically significant from the control egg chambers (unpaired T-test; P<0.0001). (K) Zoomed in images of *GR1-GAL4>UAS-GFP* and *GR1-GAL4>UAS-Lac-RNAi* stage 14 egg chambers showing the dorsal appendages (L). In the control, dorsal appendages are formed, whereas the dorsal appendages in eggs expressing *Lac-RNAi* are either deformed or absent (arrows). (M) Quantification of dorsal appendage defects from control and *SJ-RNAi* stage 14 egg chambers. *n*, total number of egg chambers measured per genotype. Data represents individual values with mean ± s.d. *P* value <0.0001. Anterior is to the left. Scale bar = 100μm.

Further examination of stage 14 SJ mutant egg chambers revealed defects in dorsal appendage morphogenesis. Dorsal appendages are tubular respiratory structures that form from two populations of the dorsal FE known as floor and roof cells (Duhart *et al.* 2017). The primary phenotypes we observed in the *SJ-RNAi*-expressing egg chambers were missing dorsal appendages, or appendages that appeared to be short or broken (Figures 3L and S2). In addition, nearly all of the *SJ-RNAi*-expressing dorsal appendages that were present appeared to have a thinner stalk than found in control egg chambers (Figure S2). In quantifying these phenotypes, both BDSC and VDRC RNAi lines against Mcr (BDSC: 52%, VDRC:18%) and Cora (BDSC: 15% and VDRC: 21%) produced egg chambers with defective dorsal appendages (Fig 3M). However, the *Nrx-IV-RNAi* BDSC line did not produce abnormal dorsal appendages, whereas 33% of VDRC *Nrx-IV-RNAi* line results in defective dorsal appendages. Moreover, 19% of *Cont-RNAi-* and 13% of *Lac-RNAi-expressing* egg chambers have either missing or broken dorsal appendages. We did not observe these phenotypes in *Sinu-RNAi*-expressing egg chambers (Figure 3M).

### SJ proteins are expressed in polar and border cells and are required for effective border cell migration

The observation that Mcr, Cont, and Nrx-IV are strongly expressed in PCs and in all FCs (Figure 1D-F), motivated us to investigate their expression during the process of border cell migration. Border cell migration occurs during stages 9-10 of oogenesis (Figure 4A). During stage 9, signaling from the anterior PCs recruits a group of 4-8 adjacent FCs to form a cluster and delaminate apically into the egg chamber. The border cell cluster is organized with a pair of PCs in the center surrounded by border cells. This cluster of cells migrates between the nurse cells until they reach the anterior side of the oocyte (Figure 4A). This process takes approximately 4-6 hours and is complete in wild type egg chambers by stage 10 of oogenesis (Prasad and Montell 2007). The border cell cluster, along with the migratory centripetal cells, collaborate to form the micropyle, a hole through which the sperm enters the egg (Figure 4A) (Montell 2003; Horne-Badovinac 2020). Previous studies show that Cora and Nrg are expressed in the PCs during their migration (Wei *et al.* 2004; Felix *et al.* 2015). To determine if other SJ proteins are also expressed during border cell migration, we stained stage 9-10 wild type egg chambers with antibodies against Mcr, Cont, Nrx-IV and Cora. To mark the PCs, we co-stained the egg chambers with Fasciclin 3 (Fas3; Snow *et al.* 1989; Khammari *et al.* 2011). Mcr, Cont, Nrx-IV and Cora are all expressed in border cell clusters throughout their migration (Figure 4B-D and Supplemental Figures 1-3). Interestingly, the expression of these SJ proteins in PCs appears highest at the interfaces between polar and border cells (Figure 4B-D and Figures S3-5). SJ protein expression is also asymmetric in the border cell cluster, with higher expression along border cells closest to the oocyte, raising the possibility that these proteins may respond to or direct leading-edge polarity in the migrating border cell cluster (red arrows in Figure 4B-D).

**Figure 4.**
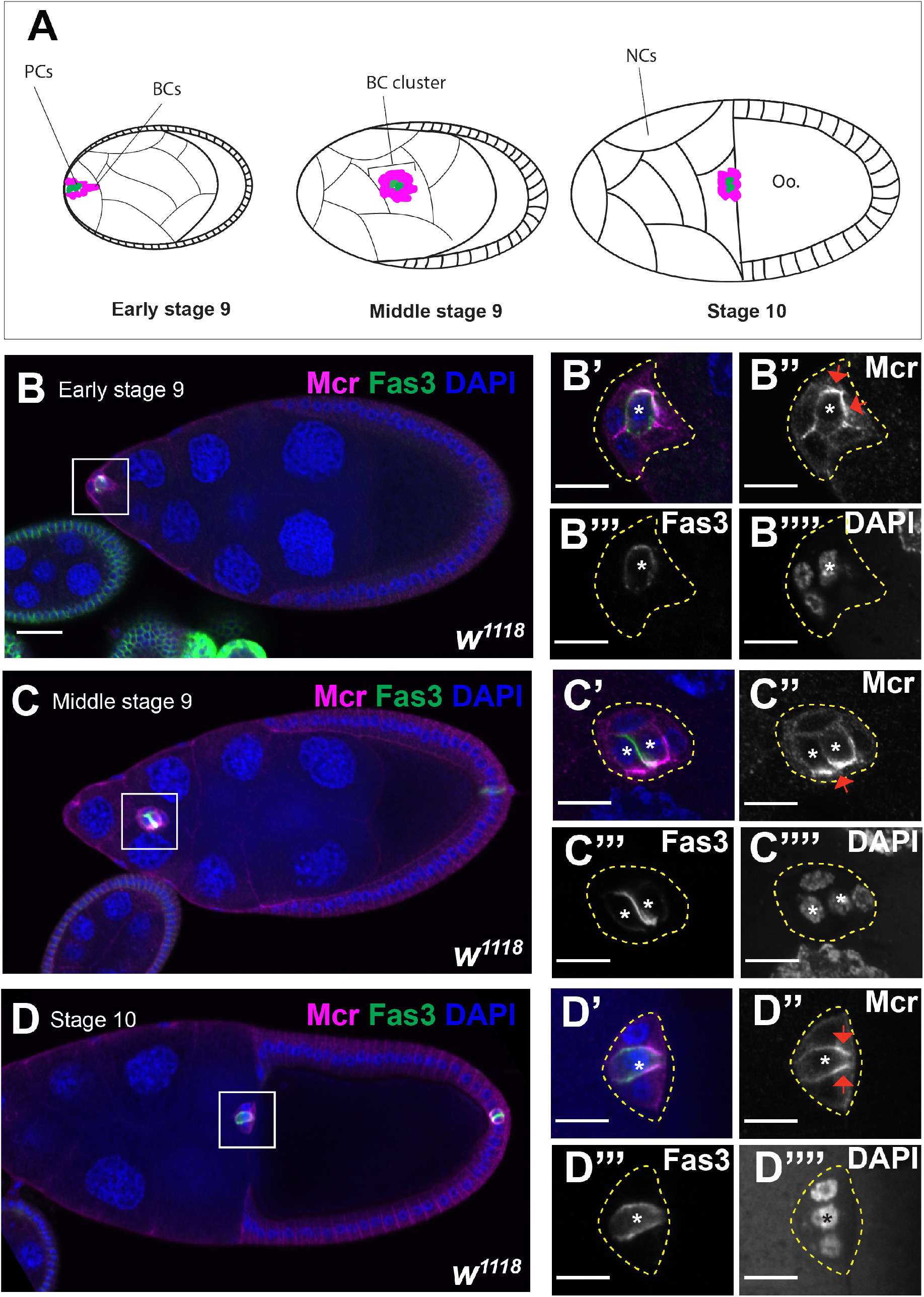
Mcr expression during border cell migration. (A) Diagram of border cell migration. At early stage 9 of oogenesis, a group of 4-6 follicle cells are specified by the polar cells (PCs) (green) to become border cells (BCs) (magenta). The BC/PC cluster delaminates apically and migrates between the nurse cells (NCs) until it reaches the oocyte (Oo.) by stage 10 of oogenesis. (B-D) Confocal optical images of wild type stage 9-10 egg chambers stained with antibodies against Mcr (magenta), Fas3 (green), and labeled with DAPI (blue). (B-D) Mcr is expressed in the PCs and BCs with the highest expression at the interface between the PCs and BCs. Mcr is enriched at the leading edge (red arrows in B”-D”). Anterior is to the left. Scale bar = 25μm in (B-D) and 10μm in (B’-B’’’’ and D’-D’’’’).

Given that Mcr, Cont, Nrx-IV and Cora are expressed in border cell clusters throughout border cell migration, we wondered if they are also required for some aspect of this process. To address this issue, we used *Slbo-GAL4* (Ogienko *et al.* 2018) to express RNAi against individual SJ genes specifically in border cells and analyzed the border cell clusters at stage 10 in these ovaries. We noticed a range of defective migration phenotypes in these egg chambers and classified them into three non-exclusive groups: failed, incomplete and dissociated clusters. Complete migration (Figure 5A-C) is characterized by having the entire border cell cluster physically touching or just adjacent the oocyte by the end of stage 10 (Figure 5C). A failed cluster is characterized by a border cell cluster that has not delaminated from the FE by stage 10 (Figure 5D). An incomplete migration phenotype is characterized as an intact cluster that has not reached the oocyte by the end of stage 10 (Figure 5E, where the two dashed lines indicate the range of distances at which border cell clusters were categorized as incomplete). Finally, a dissociated cluster phenotype is characterized by a cluster that has broken into a linear string of border cells or that has one or more border cells that remain between nurse cells and are not connected to the larger border cell cluster (Figure 5F and H).

**Figure 5.**
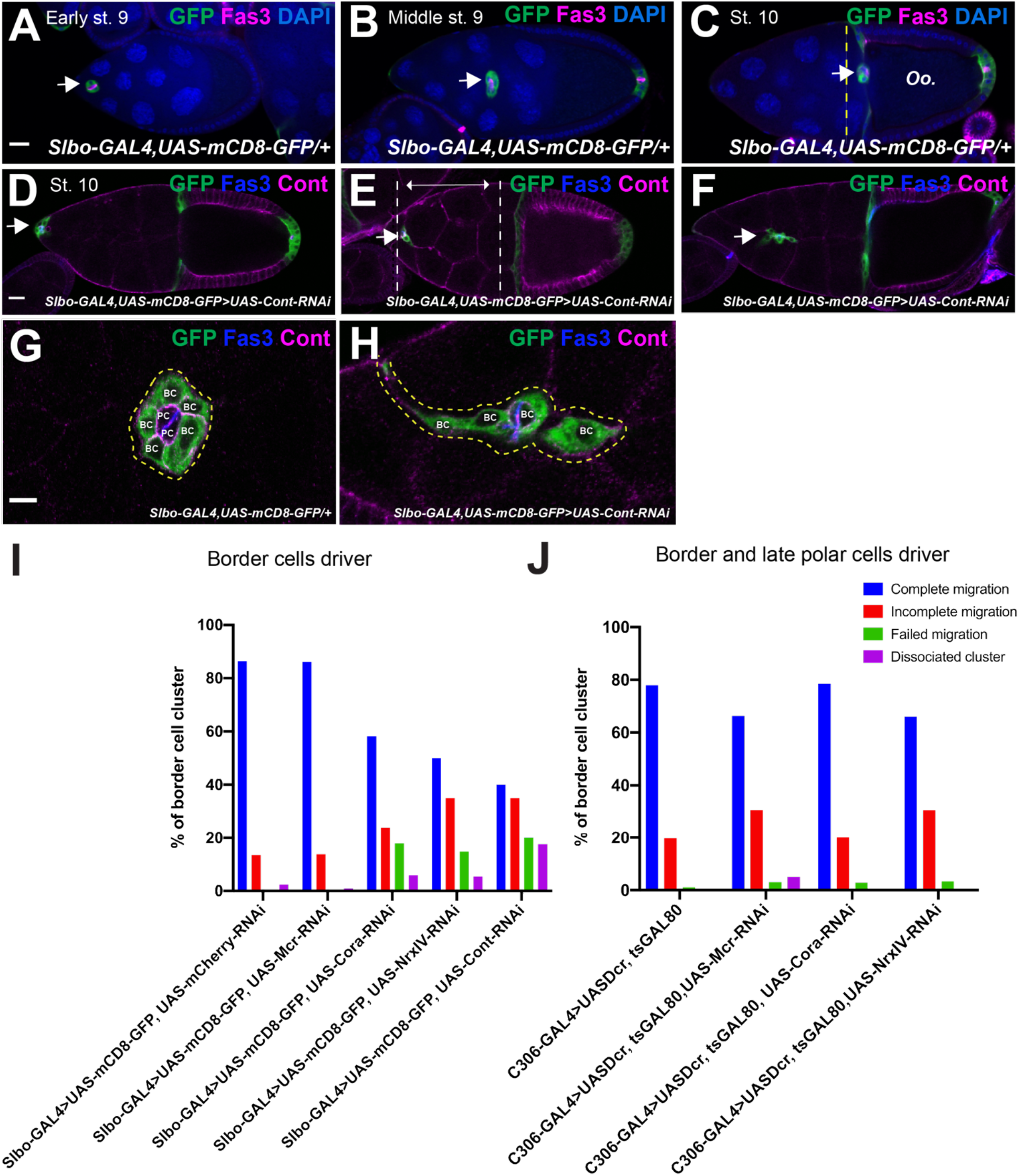
Mcr, Cont, Nrx-IV, and Cora are required for effective border cell migration. (A-C) Border cell migration in control egg chambers. Egg chambers are immunostained with anti-Fas3 (magenta) to mark the polar cells, GFP (green) to mark border cells and labeled with DAPI (blue). Arrow indicates a border cell cluster that has completed migration to the oocyte by stage 10. (D-F) Stage 10 egg chambers expressing *Cont-RNAi* in border cells immunostained with anti-Cont (magenta) and anti-Fas3 (blue) as examples of BC cluster migration defects in *SJ-RNAi* egg chambers. Failed (arrow in D), incomplete (arrow in E), or disassociated border cell migration (arrow in F). Higher magnification of wild type (G) and Cont-RNAi-expressing (H) border cell cluster showing the dissociation of border cell cluster observed in *SJ-RNAi* clusters. (I and J) Quantification of border cell cluster phenotypes at stage 10 egg chambers in control and *SJ-RNAi* driven by either Slbo-GAL4 (I) or C306-GAL4 (J) line. Anterior is to the left. Scale bar = 25μm.

In control stage 10 egg chambers (*Slbo-GAL4; UAS-mCD8-GFP/UAS-mCherry-RNAi*), 86% (n=81) of border cell clusters completed their migration, with the remainder showing incomplete migration (Figure 5I). In contrast, stage 10 egg chambers expressing RNAi against *Cora, Nrx-IV*, or *Cont* in the border cells resulted in 58% (n=67), 50% (n=74), and 40% (n=85) of border cell clusters completing migration, respectively (Figure 5I). The remaining *Cora-RNAi-* and *Nrx-IV-RNAi-*border cell clusters showed a combination of incomplete migration or failed to delaminate (Figure 5I). *Cont-RNAi-border* cell clusters also showed 35% of incomplete border cell migration, but additionally had 17% of the clusters disassociating during their migration (Figure 5E, F, H and I). *Mcr-RNAi-border* cell clusters were indistinguishable from controls with 86% (n=94) completing their migration and the remainder showing only 13% incomplete migration (Figure 5I).

To extend these studies, we examined border cell migration in egg chambers expressing RNAi against SJ genes using the *C306-GAL4* driver. *C306-GAL4* is expressed in the border cells throughout the process of border cell migration (Murphy and Montell 1996) and in PCs just at stage 10 (H.A., unpublished observation). In control egg chambers (*C306-GAL4/UAS-Dcr; GAL80^ts^/+*), 78% (n=91) of BC clusters completed their migration and 19% displayed incomplete migration (Figure 5J). Stage 10 egg chambers expressing *C306>Mcr-RNAi* displayed 66% (n=98) complete border cell migration with 30% showing incomplete migration, 5% showing dissociated clusters and 3% showing failed border cell migration (Figure 5J). Similarly, egg chambers expressing *C306>Nrx-IV-RNAi* displayed 66% (n=59) complete border cell migration with 30% showing incomplete migration and 3% failing to delaminate (Figure 5J). Finally, 78% (n=70) of stage 10 *C306>cora-RNAi-expressing* border cells displayed complete border cell migration, whereas 20% showed incomplete migration and 3% failed to delaminate (Figure 5J).

### SJ biogenesis in the follicular epithelium

The redistribution of SJ proteins in the FCs of later stage egg chambers resembles the dynamic relocalization of SJ proteins during the biogenesis of the junction during embryogenesis (Tiklová *et al.* 2010). In embryonic epithelia, SJ protein enrichment at the apical-lateral domain (presumptive SJ) requires endocytosis and recycling of SJ proteins to the membrane, and is interdependent on the presence of all core SJ proteins (Ward *et al.* 1998; Hall *et al.* 2014). Coincident with the strong localization of SJ proteins to the presumptive SJ at stage 16 of embryogenesis, ladder-like electron-dense intermembrane septae are visible by electron microscopy that become progressively organized by stage 17 (Schulte *et al.* 2003; Hildebrandt *et al.* 2015). We therefore sought to determine if similar processes occur during the formation of SJs in the FE.

To test for the interdependence of SJ proteins for localization, we examined Cora and Mcr expression in wild type, *Mcr-RNAi*, and *Nrx-IV-RNAi* stage 12 FCs. In wild type stage 12 egg chambers, Cora is strongly co-localized with Mcr at the apical-lateral domain of the FCs (completely penetrant in 95 cells from 31 egg chambers) (Figure 6A). In contrast, Cora and Mcr are mislocalized along the lateral domain in stage 12 *Nrx-IV-RNAi* FCs (Figure 6B), much like they are in stage 2-8 wild type FCs (Figure 1D). Specifically, 6 out of 20 *Nrx-IV-RNAi* FCs cells from 19 egg chambers showed complete mislocalization, whereas 13 of these 20 cells showed largely lateral localization with some degree of apical enrichment. Similarly, in stage 12 *Mcr-RNAi* FCs, Cora was mislocalized along the lateral membrane in 39 out of 47 cells examined from 19 egg chambers, with the remaining 8 cells showing some enrichment of Cora at the apical lateral domain (Figure 6C). Notably, cells that showed apical enrichment of Cora also retained small foci of Mcr expression that overlaps with the enriched Cora (Figure 6D), suggesting the perdurance of Mcr in these cells may have allowed for normal SJ organization. Together, these experiments indicate that SJ biogenesis in the FE of late-stage egg chambers requires the expression of at least some core SJ proteins.

**Figure 6.**
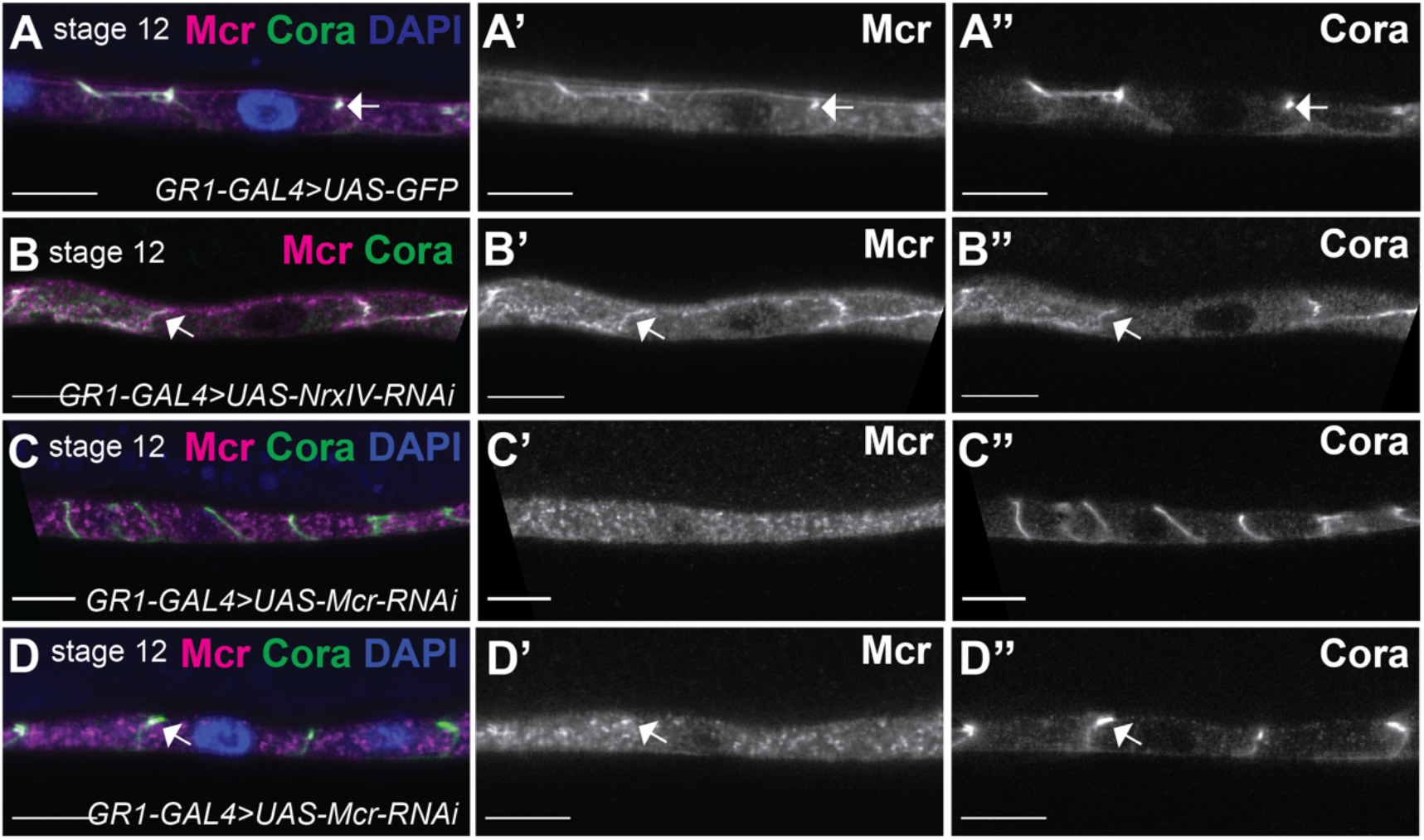
The apical-lateral localization of Cora depends on Mcr localization at the apical-lateral membrane. (A-D) Confocal optical sections of stage 12 follicle cells (FCs) stained with Mcr (magenta) and Cora (green) and labeled with DAPI (blue). Mcr and Cora co-localize at the septate junction (SJ) (A), whereas Mcr and Cora localize along the lateral membrane of *NrxIV-RNAi* expressing FCs (B). In *Mcr-RNAi* expressing FCs, Cora localizes laterally (C) or maintains its enrichment at the apical-lateral membrane (D). Scale bar = 10μm (A-D) and 0.5μm (E and F).

We next wanted to investigate whether the relocalization of SJ proteins to the presumptive SJ required endocytosis and recycling. In the embryonic hindgut, Cora, Gliotactin (Gli), Sinu, and Melanotransferrin (Mtf) localize with the early endosomal marker, Rab5, and partially localize with the recycling marker, Rab11 during SJ biogenesis (Tiklová *et al.* 2010). Moreover, blocking Rab5 or Rab11 function prevents Cora, Gli, Sinu, and Mtf apical-lateral localization (Tiklová *et al.* 2010), and thus SJ formation. To determine if similar processes occur during SJ formation in FCs, we expressed a dominant negative version of Rab5 (*UAS-Rab5^DN^*) in FCs using *GR1-GAL4* and examined the expression of Mcr and Cora in stage 11 FCs. In wild-type FCs at that stage, Mcr and Cora are enriched at the apical-lateral membrane (completely penetrant in 91 cells from 15 egg chambers; arrows in Figure 7A), whereas both Mcr and Cora remains localized along the lateral membrane in the Rab5^DN^-expressing FCs (n=97 cells, 15 egg chambers; Figure 7B). Similarly, Cora and Mcr co-localize at the apical-lateral membrane of the FCs of stage 12 egg chambers (n=64 cells, 15 egg chambers; arrows in Figure 7C), whereas knocking down the expression of Rab11 in stage 12 FCs results in the mislocalization of Cora and Mcr (n=23 cells out of 44, 16 egg chambers; arrow in Figure 7D). Cora is exclusively mislocalized along the lateral membrane in these cells, whereas Mcr is most frequently missing from the plasma membrane, either in punctate cytoplasmic foci or completely gone (in 21 of the 23 *Rab11-RNAi* cells; Figure 7D). Interestingly, we noted that the FE in *Rab5^DN^-* and *Rab11-RNAi*-expressing egg chambers are taller in the apical/basal dimension than similarly staged wild type egg chambers (compare Figure 7A with 7B and Figure 7C with 7D), although the effect is greater with *Rab5^DN^* than with *Rab11-RNAi.* Taken together, these results suggest that similar to embryonic epithelia, the maturation of SJs in the FE requires Rab5-mediated endocytosis and Rab11-mediated recycling.

**Figure 7.**
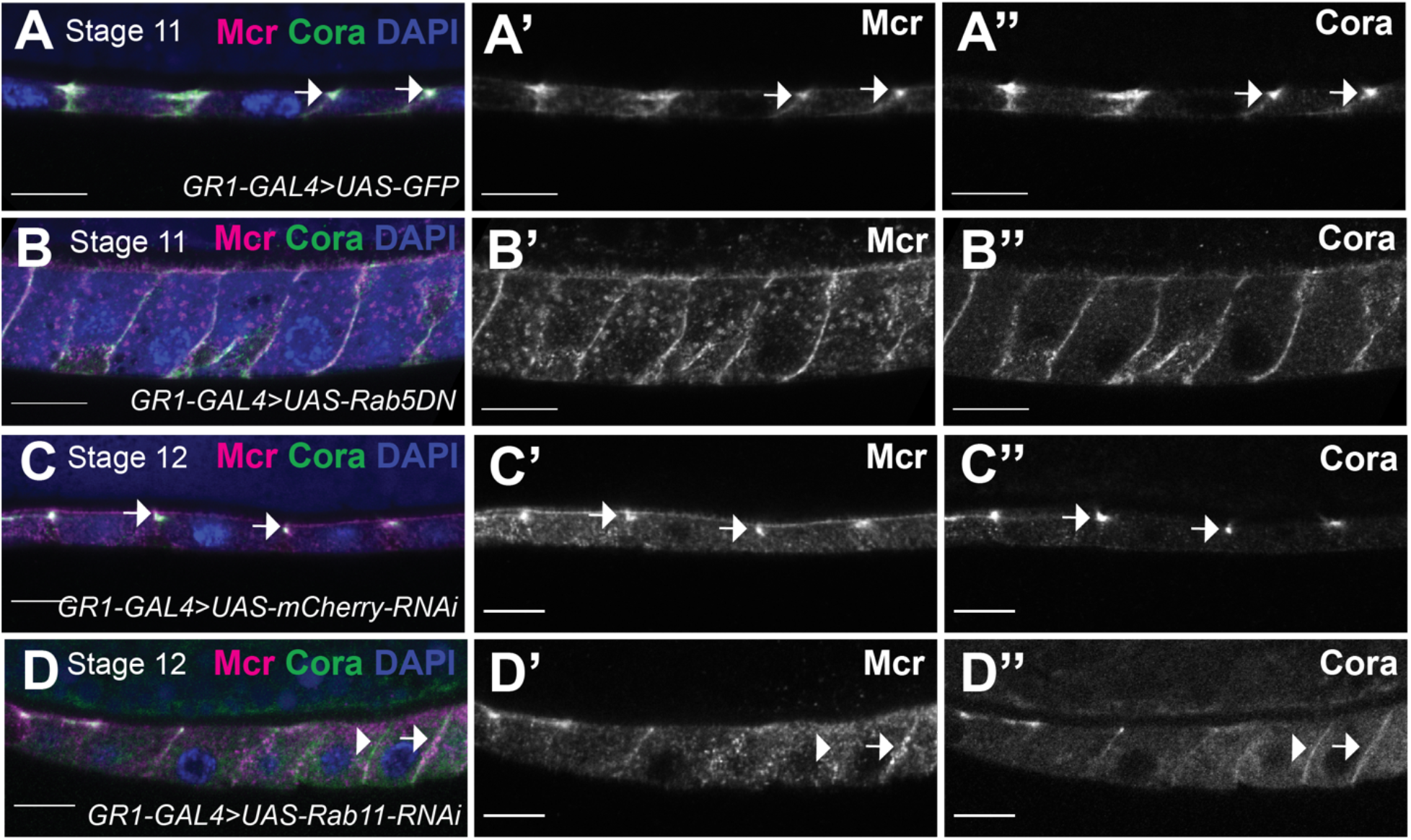
Mcr and Cora require Rab5 and Rab11 for their correct localization at the SJ. While Mcr and Cora co-localize at the apical-lateral membrane of stage 11 FCs (arrows in A-A’’), both Cora and Mcr fail to localize at the SJ in Rab5^DN^ expressing FCs (B). In stage 12 FCs, Mcr and Cora co-localize at the apical-lateral membrane (C). (D) In *Rab11-RNAi* expressing FCs, however, Cora localizes along the lateral membrane, whereas Mcr is either missing (arrowhead) or localizes along the lateral membrane (arrow). Anterior is to the left. Scale bars = 10μm.

## Discussion

In this study, we have demonstrated that a subset of SJ proteins is expressed in egg chambers throughout oogenesis and are required for critical morphogenetic processes that shape the egg, including egg chamber elongation, dorsal appendage formation and border cell migration (required to form the micropyle). Interestingly, the subcellular localization of these SJ proteins in the ovarian FCs changes coincident with the establishment of the occluding junction in much the same way that they do during embryogenesis in ectodermal epithelial cells (Tiklová *et al.* 2010), suggesting that a similar maturation process for the SJ occurs in this tissue.

### Biogenesis of the SJ in the FE

The mechanisms of SJ biogenesis in embryonic epithelia has been well-studied and involves several steps in which transmembrane SJ proteins are first localized all along lateral plasma membranes (by stage 12 of embryogenesis), but then must be endocytosed and recycled back to the plasma membrane prior to aggregating in the region of the presumptive SJ (between stages 13 and 16; Tiklová *et al.* 2010). Prior to this relocalization step, SJ proteins show high mobility in the plane of the membrane by Fluorescence Recovery After Photobleaching (FRAP) analysis, but become strongly immobile as the relocalization is occurring (Oshima and Fehon 2011). As these steps are occurring (e.g. stage 14 of embryogenesis), electron-dense material begins to accumulate between adjacent cells in the presumptive SJ that takes on the appearance of ladder-like septae by stage 17 of embryogenesis (Tepass and Hartenstein 1994). Functional studies indicate that the occluding function of the junction is established late in embryogenesis, near the end of stage 15 (Paul *et al.* 2003). Importantly, the process of SJ biogenesis is interdependent on the function of all core components of the junction and several accessory proteins including Rab 5 and Rab 11 (Ward *et al.* 1998; Genova and Fehon 2003; Tiklová *et al.* 2010).

Here, we observe that many of these same steps occur in the maturation of SJs in the FE. We first show that SJ proteins are expressed in the FE beginning early in ovarian development where they localize all along the lateral membrane (Figure 1). These proteins appear to become enriched at the presumptive SJ by stage 11 (Figure 6). The relocalization of SJ proteins to the SJ requires core SJ proteins including Mcr and Nrx-IV, and accessory proteins including Rab 5 and Rab 11 (Figures 6 and 7). Prior studies indicate the presence of electron dense material between FE cells as early as stage 6 (Müller 2000), with a ladder-like appearance of extracellular septae by stage 10B (Mahowald 1972). A recent study of the occluding function in the FE show a similar pattern of protein localization for endogenously tagged Neuroglian-YFP and Lachesin-GFP, and demonstrates that an intact occluding junction is formed during stage 11 of oogenesis (Isasti-Sanchez *et al.* 2020). It is interesting to note the FE is a secondary epithelium initiated by a mesenchymal to epithelial transition (Tepass *et al.* 2001), and yet SJ biogenesis appears to be very similar to that observed in the primary epithelia found in the embryo.

### SJ proteins are required for morphogenetic events during oogenesis

The similarities in the dynamic expression of SJ proteins in the FE and embryonic epithelia, coupled with the observation that SJ proteins are required for numerous embryonic developmental events (Hall et al. 2014) and references therein) motivated us to explore whether SJ proteins have similar roles in morphogenetic processes that shape the egg. Using a targeted RNAi approach, we show that reducing the expression of Mcr, Nrx-IV, Cont, Cora, Sinu, or Lac throughout oogenesis in the FCs results in significantly rounder stage 14 egg chambers, with many showing additional defects in dorsal appendages (Figures 3 and S2). The initiation and maintenance of egg elongation is achieved at various stages throughout oogenesis (Gates 2012; Bilder and Haigo 2012; Cetera and Horne-Badovinac 2015). From stage 3-6, a gradient of JAK-STAT signaling is required at each pole of the egg chamber to promote Myosin II-based apical cell contractions (Alégot *et al.* 2018). From stage 6-8, the formation of a robust planar polarized molecular corset - consisting of basal actin cytoskeleton and basement membrane protein fibrils - is required for egg elongation, and requires collective FC migration over a static basement membrane (Gutzeit *et al.* 1991; Frydman and Spradling 2001; Bateman *et al.* 2001; Viktorinová *et al.* 2009; Haigo and Bilder 2011; Horne-Badovinac *et al.* 2012; Cetera *et al.* 2014; Isabella and Horne-Badovinac 2016; Campos *et al.* 2020). During the middle stages of oogenesis (stages 9 and 10), basal actin stress fibers undergo actomyosin contractions, which contribute to egg elongation (He *et al.* 2010; Qin *et al.* 2017). Finally, later in oogenesis, the maintenance of a planar polarized molecular corset is required to retain an elongated egg chamber shape (Haigo and Bilder 2011; Cha *et al.* 2017; Campos *et al.* 2020). Future studies are needed to determine if SJ proteins are required for the establishment and/or maintenance of egg shape. Since many of the events involved in egg elongation occur prior to the formation of a functional (and ultrastructurally intact) occluding junction, it raises the possibility that SJ proteins have a function in morphogenesis that is independent of their role in forming a tight occluding junction, much as they do in the embryo.

Stage 14 egg chambers from many of the *SJ-RNAi* lines possessed aberrant dorsal appendages, often characterized by misshapen, broken or missing appendages (Figures 3 and S2). The formation of dorsal appendages occurs during stages 10B-14 and requires cell shape changes and cell rearrangements, coupled with chorion protein secretion (Dorman et al. 2004). Similar morphogenetic processes are required for dorsal closure and head involution during embryogenesis (Hayes and Solon 2017; VanHook and Letsou 2008), defects strongly associated with zygotic loss of SJ expression in the embryo (Hall and Ward 2016). We are interested to determine if the mechanism by which SJ proteins contribute to dorsal appendages formation and dorsal closure and head involution are similar. Potential roles could involve regulating the cytoskeleton to facilitate cell shape changes and rearrangements, but another intriguing possibility is that SJ proteins may also be required for chorion protein secretion or crosslinking. Broken and missing dorsal appendages may result from mechanical disruption due to chorion defects. We also noticed mature *SJ-RNAi* eggs with a thin chorion (data not shown). Notably, embryos with mutations in many different SJ genes show defects in the embryonic cuticle including faint denticle belts and delamination of cuticle layers (Lamb *et al.* 1998; Hall and Ward 2016).

Our observation that specifically knocking down the expression of several SJ proteins in border cells results in defective border cell migration (Figure 5) supports a role for SJ proteins in morphogenesis, independent of their role in forming an occluding junction. The phenotypes we observed include failing to complete migration and partial disassociation of the complex by the end of stage 10, which is prior to the formation of an intact SJ. Although the penetrance of these phenotypes is mild (Figure 5I and J), it is possible that these phenotypes underestimate the full requirement of SJ proteins in border cell migration since this process takes a relatively short time (4-6 hours) (Prasad and Montell 2007), while SJ proteins are thought to be very stable in the membrane (Oshima and Fehon 2011). One caveat to the idea that perdurance may account for the mild phenotypes is that *C306-GAL4* does not appear to produce a stronger phenotype than *slbo-GAL4*, even though it is expressed earlier and is presumably knocking down SJ proteins longer. Perhaps knocking the proteins down more quickly using the DeGradFP system (Caussinus, Kanca, and Affolter 2012) could address this possibility in the future. These phenotypes also indicate a potential role for SJ proteins in cell adhesion and/or cell polarity during migration. Specifically, we note that Mcr appears to be enriched in polar cell membranes that contact border cells at the leading edge of the cluster (those that are oriented closest to the oocyte) in wild type egg chambers (Figure 4). Whether SJ proteins are required for aspects of planar polarity in the border cell cluster is an interesting unanswered question. Perhaps the incomplete migration defect results from a meandering migration through the nurse cells, something that has been observed for knockdown of DE-Cadherin in border cells (Cai et al. 2014). Live imaging studies should be able to distinguish between pathfinding defects and a general reduction in speed or premature stopping. A role for SJ proteins in cell adhesion in the ovary has been reported previously. Reducing the level of Nrg in FCs results in the failure of newly divided FCs to reintegrate into the FE, indicating a role for *Nrg* in lateral cell adhesion (Bergstralh *et al.* 2015). In addition, expressing a null allele of *Nrg* in FCs enhances the invasive tumor phenotype of a *Discs Large (Dlg*) mutation (Wei *et al.* 2004).

## Acknowledgements

We thank Jocelyn McDonald, the Bloomington *Drosophila* Stock Center, and the Vienna *Drosophila* RNAi Center for fly stocks. We thank Manzoor Bhat and the Developmental Studies Hybridoma Bank (created by the NICHD of the NIH and maintained at The University of Iowa, Department of Biology, Iowa City, IA 52242) for antibodies used in this study. We thank Brian Ackley for the use of his Olympus FV1000 confocal microscope. We thank Lindsay Ussher for preliminary studies on border cell migration and thoughtful discussions on the project. We also thank Sally Horne-Badovinac, Jocelyn McDonald, Yujun Chen, and members of the Ward lab for helpful discussion about the project and manuscript.

