## supplemental figures for "Septate junction proteins are required for egg elongation and border cell migration during oogenesis in Drosophila"

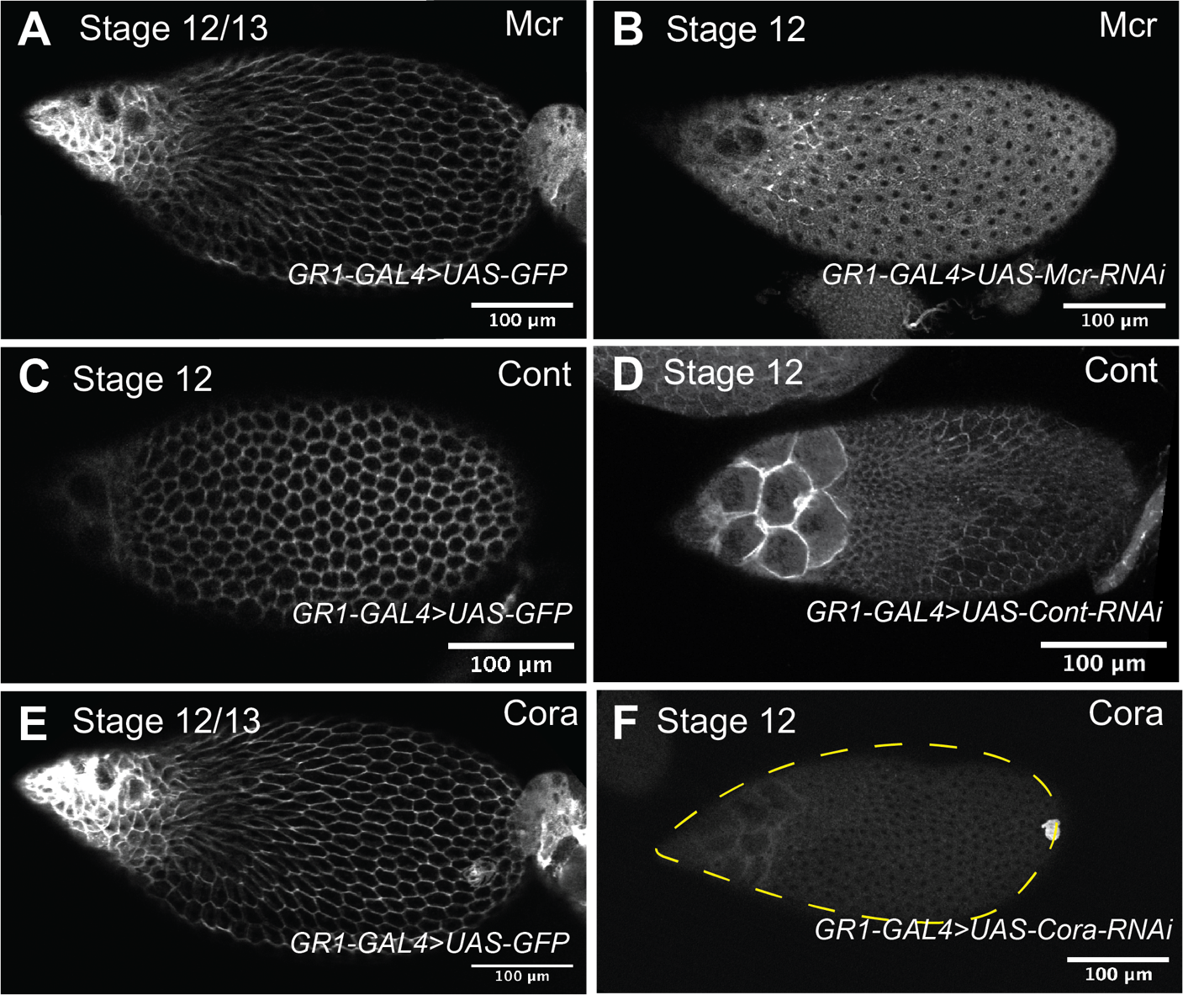

**Figure S1. Mcr, Cont, and Cora expression in *GR1>SJ-RNAi* follicle cells**. (A, C, and E) Confocal sections of stage 12/13 (A and E) and stage 12 (C) control (*GR1>GFP*) egg chambers stained with Mcr (A), Cont (C), and Cora (E). (B, D, and F) Confocal sections of stage 12 egg chambers expressing *GR1>Mcr-RNAi* (B), *GR1>Cont-RNAi* (D), or *GR1>Cora-RNAi* (F) stained with antibodies against Mcr (B), Cont (D) or Cora (E). Note that level of the proteins is significantly knocked down in the *RNAi-*expressing egg chambers. A and E are images from the same egg chamber co-stained for Mcr (A) and Cora (E). Anterior is to the left and posterior is to the right. Scale bar = 100µm.

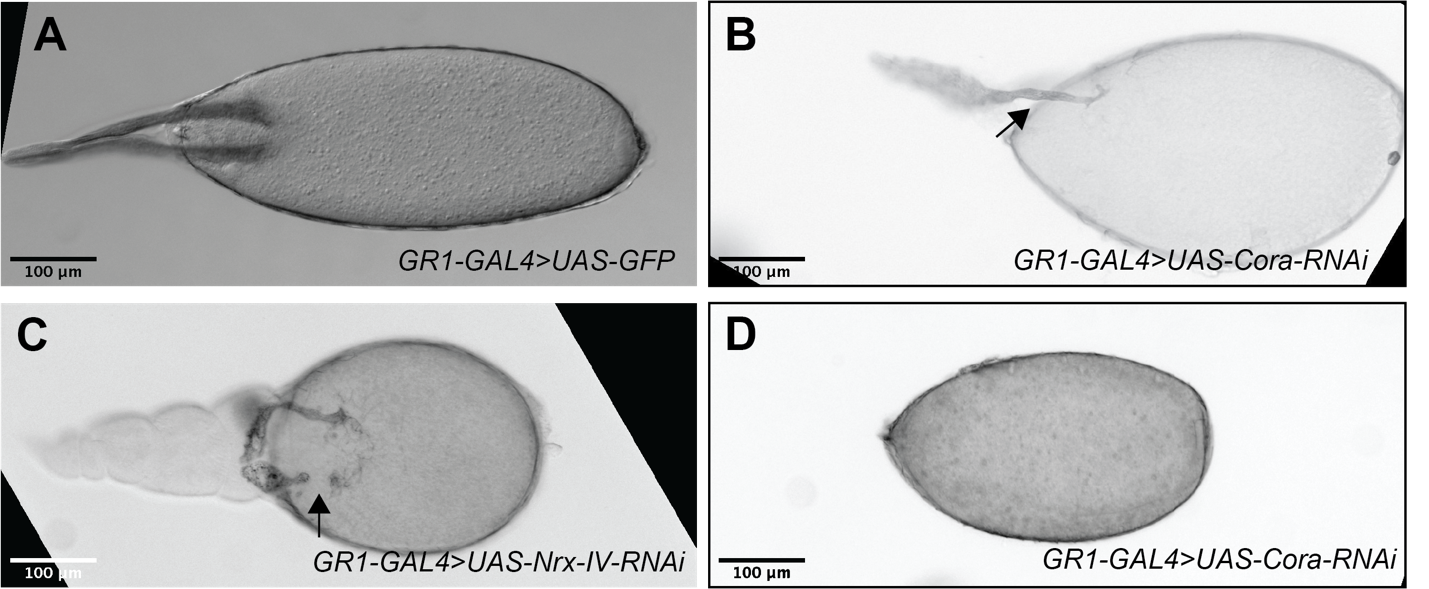

**Figure S2. Examples of dorsal appendages defects in SJ-RNAi stage 14 egg chambers**. (A–D) Brightfield photomicrographs of control *GR1>GFP* (A), *GR1>Cora-RNAi* (B and D) and *GR1>Nrx-IV-RNAi* (C) stage 14 egg chambers. Dorsal appendages in control egg chambers show uniform thickness along the stalk of the appendage that becomes thicker and broader at the paddle (A). In contrast the shape of dorsal appendages in SJ-RNAi egg chambers show a range of abnormal phenotypes including thinner stalks, particularly near the oocyte (black arrow in B), with wider and flatter paddles, and broken (black arrow in C) or missing dorsal appendages (D). Note that the spacing between appendages can also be larger in SJ-RNAi egg chambers (C). Anterior is to the left and posterior is to the right. Scale bar = 100µm.

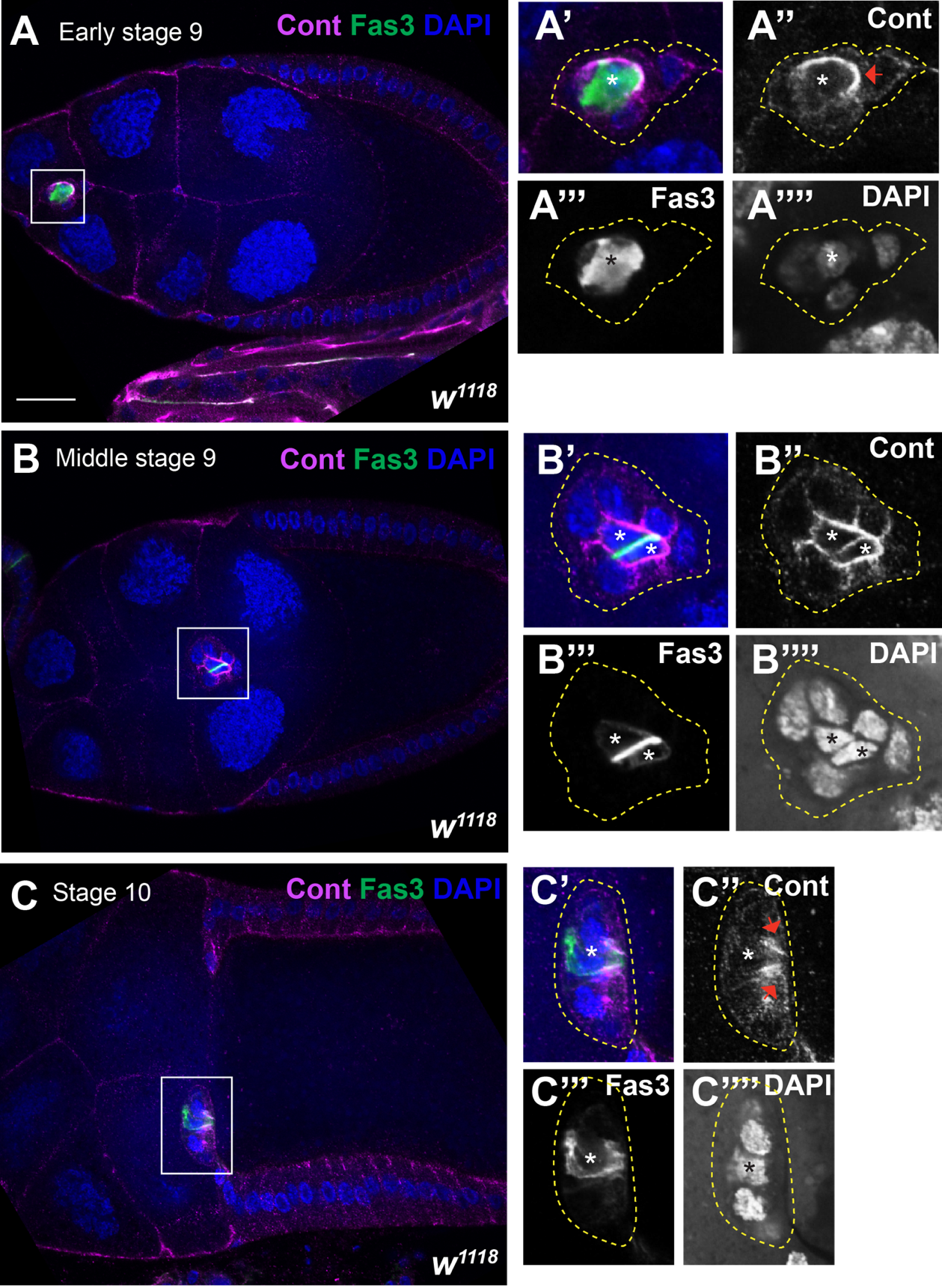

**Figure S3. Cont expression in the border cluster throughout border cell migration.** (A–C) Confocal sections of wild type early stage 9 (A), middle stage 9 (B), and stage 10 (C) stained with Cont (magenta), Fas3 to mark the polar cells (green) and labeled with DAPI (blue). (A’–A’’’’, B’–B’’’’, and C’–C’’’’) are a zoomed in images of the border cell cluster in A-C. Cont is expressed in the border cell cluster with the strongest expression in the polar cells. Cont also shows a symmetric localization at the leading edge of the cluster (red arrows in A’’ and C’’). Anterior is to the left. Scale bar= 25µm.
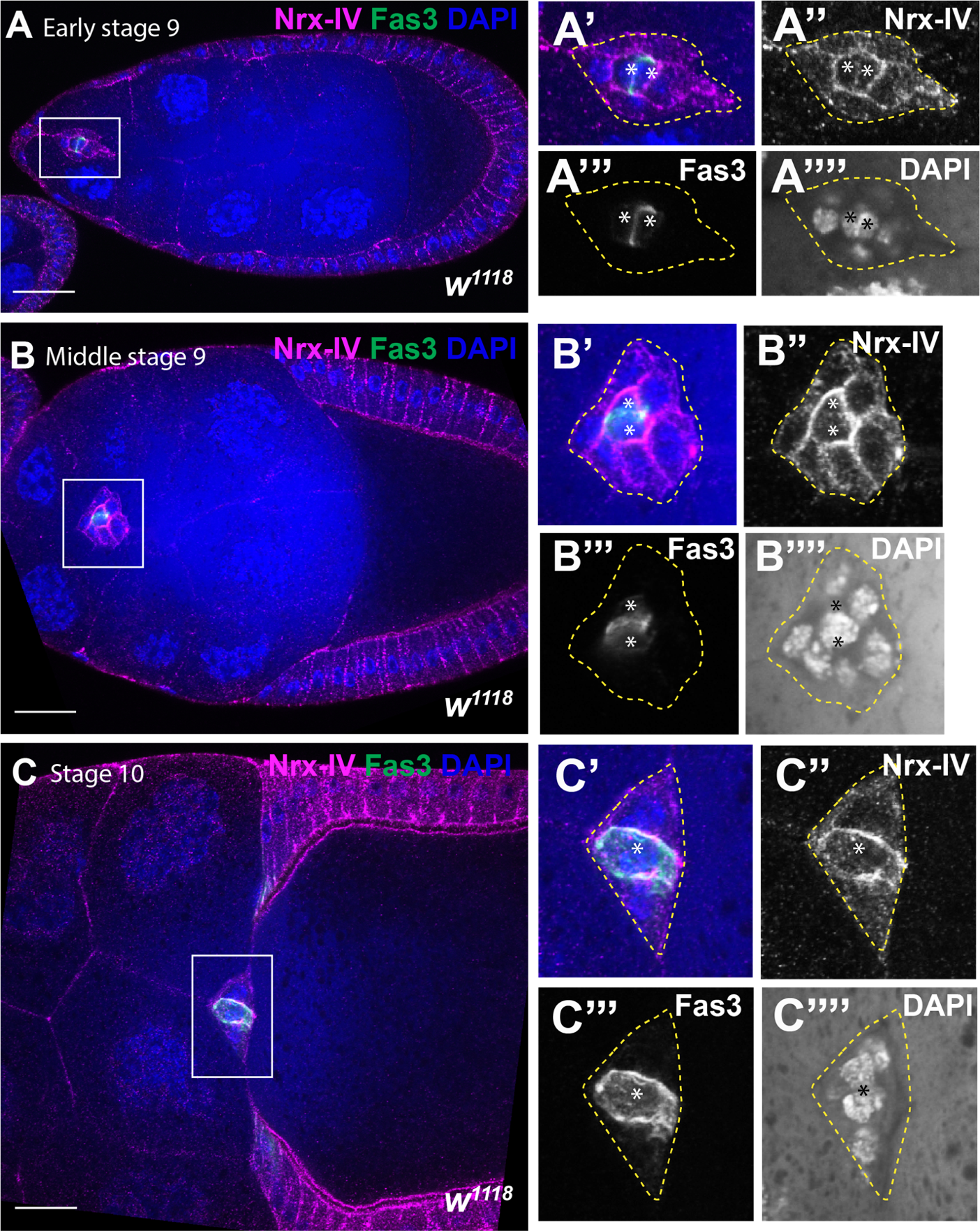

**Figure S4. Nrx-IV expression in the border cell cluster during border cell migration.** (A–C) Confocal sections of early stage 9 (A), middle stage 9 (B), and stage 10 wild type egg chambers stained with Nrx-IV (magenta), Fas3 to mark the polar cells (green) and labeled with DAPI (blue). (A’–A’’’’), (B’-–B’’’’), and (C–C’’’’) are zoomed in images of A, B, and C. Note the localization of Nrx-IV in the border cell cluster throughout border cell migration with enrichment at the PCs-BCs contacts (A’’, B’’, and C’’). Anterior is to the left. Scale bar= 25µm.

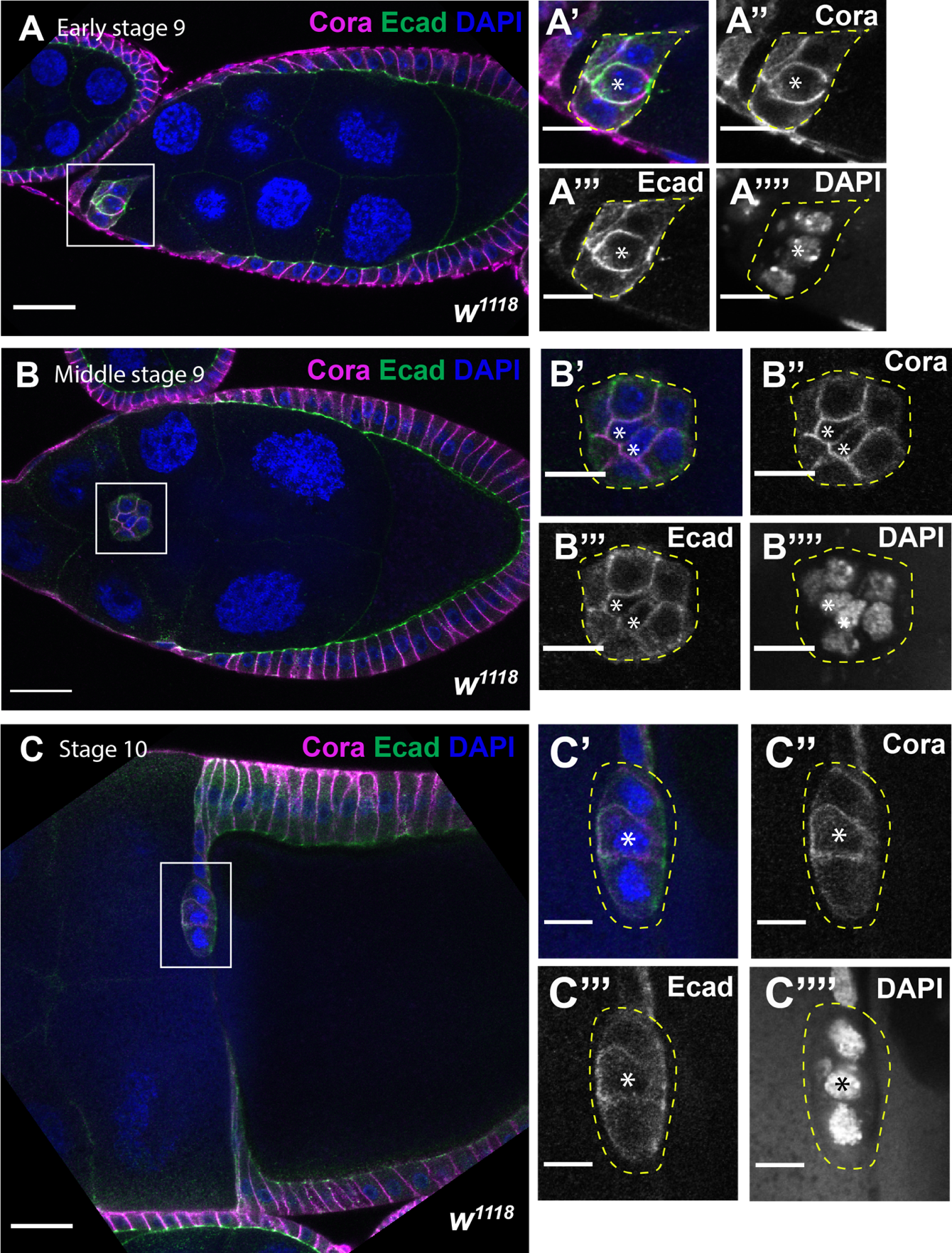

**Figure S5. Cora expression in the border cell cluster during border cell migration.** (A–C) Confocal sections of wild-type egg chambers at early stage 9 (A), middle stage 9 (B), and stage 10 (C) stained with Cora (magenta), Ecad (green), and labeled with DAPI (blue). (A’–A’’’’), (B’–B’’’’), and (C’–C’’’’) are zoomed in images of A, B, and C. Cora is expressed in the polar cells and border cells throughout border cell migration with high level at the polar cells-border cells interface (A’’, B’’, and C’’). Anterior is to the left. Scale bar= 25µm.
